# Emodin-Enhanced hUC-MSC Extracellular Vesicles Alleviate Acute Pancreatitis by Targeting Inflammation and Pyroptosis

**DOI:** 10.1101/2024.10.22.619584

**Authors:** Xiong Liu, Xianwei Huang, Xiaodong Huang, Siyao Liu, Jun Hu, Lin Jiyan

**Affiliations:** Department of Emergency, the First Affiliated Hospital of Xiamen University, School of Medicine, Xiamen University, Xiamen, China; Xiamen Key Laboratory for Clinical Efficacy and Evidence-Based Research of Traditional Chinese Medicine, Xiamen, China

**Keywords:** Extracellular vesicle, Human umbilical cord mesenchymal stem cells (hUC-MSCs), Acute pancreatitis, Emodin, AR42J cells, Pyroptosis

## Abstract

Acute pancreatitis (AP) is a complex condition requiring immediate treatment. Both extracellular vesicles derived from human umbilical cord mesenchymal stem cells (hUC-MSC-EVs) and emodin, a naturally occurring anthraquinone used in traditional Chinese medicine, have shown therapeutic potential in treating AP. However, the mechanisms by which hUC-MSC-EVs and emodin alleviate AP, and whether they exert a synergistic effect on inflamed pancreatic tissues, remain unclear.

In this study, we developed AP cell, organoid, and animal models to compare the effects of emodin, hUC-MSC-EVs, and emodin-loaded hUC-MSC-EVs on cell viability, inflammation, and pyroptosis. Our data revealed that all three treatments improved cell viability, reduced pro-inflammatory cytokine expression, and inhibited pyroptosis in the AP models. Notably, the encapsulation of emodin significantly enhanced the protective effects of hUC-MSC-EVs.

These findings suggest that emodin’s protective effects on inflamed pancreatic tissues may be attributed, at least in part, to its anti-inflammatory and anti-pyroptotic properties. Additionally, our study proposes a novel strategy for engineering hUC-MSC-EVs for potential therapeutic applications in AP treatment.

## Introduction

The pancreas is a vital organ responsible for secreting digestive enzymes. Pancreatitis, or inflammation of the pancreas, manifests with symptoms such as nausea, vomiting, severe upper abdominal pain, fever, hypotension, and can even lead to organ failure. Along with common risk factors like gallstones, alcoholism, hyperlipidemia, and iatrogenic causes such as ERCP/EUS, pancreatitis can also result from metabolic, genetic, infectious, and lifestyle factors(1). Around 20% of acute pancreatitis (AP) patients develop a moderately severe or severe form, which may involve organ failure, pancreatic necrosis, the need for intensive care, or interventions for complications like necrosectomy. The overall mortality rate for AP is approximately 2%, but in severe or necrotizing cases, it can reach up to 20%(2). Current treatments focus on fluid resuscitation, nutritional support, analgesia, endoscopic intervention, and surgery(3); but no drugs specifically target the inflamed pancreas(4). Thus, developing new treatments to reduce mortality in severe pancreatitis is crucial.

AP is characterized by the breakdown of pancreatic tissue and activation of a systemic inflammatory response. This process is primarily driven by the premature activation of trypsinogen into trypsin within acinar cells. In addition, early injury responses in the pancreatic duct and endothelial cells further contribute to the initiation and perpetuation of the disease(5). The premature activation leads to tissue injury and the release of damage-associated molecular patterns (DAMPs), which triggers neutrophil recruitment and a subsequent inflammatory cascade(6-8).

Pyroptosis, a form of programmed cell death, is marked by cell swelling, inflammasome formation, membrane rupture, and the release of inflammatory cytokines such as IL-18. This process is distinct from other types of cell death, such as apoptosis, necrosis, and ferroptosis (9, 10). Initially thought to be a defense mechanism for immune cells against pathogens, pyroptosis is now recognized as playing a key role in various physiological and pathological processes (11, 12). In AP, pyroptosis contributes to the disease by promoting the release of DAMPs(13, 14). Therefore, targeting pyroptosis presents a potential strategy for AP treatment. For example, IL-37 can alleviate AP by inhibiting pyroptosis in pancreatic acinar cells (15), and blocking the NLRP3 inflammasome can reduce inflammation in AP models (16).

Traditional Chinese medicine has demonstrated significant therapeutic effects in treating AP (17). Emodin, a compound derived from the rhizomes and roots of plants like Rheum palmatum, possesses anti-inflammatory, antiviral, antibacterial, and immunomodulatory properties and has been effective in treating various diseases, including AP(17). Mechanistically, emodin inhibits NLRP3 inflammasome activation and pyroptosis through the Nrf2/HO-1 signaling pathway, thereby alleviating AP-induced lung injury(18). It also prevents the progression of AP by modulating lncRNA TUG1(19). However, the precise mechanisms of emodin’s action, particularly its direct effects on inflamed pancreatic cells, remain unclear.

Stem cell-based therapies have been developed and tested for various medical conditions, including AP. The success of these treatments relies on several factors, especially the availability of safe and easily accessible stem cells, such as adult stem cells and induced pluripotent stem cells (iPSCs) (20). Human umbilical cord mesenchymal stem cells (hUC-MSCs) are particularly promising for clinical use due to their advantageous properties, including ease of ex vivo expansion and noninvasive collection methods(21, 22). Importantly, hUC-MSCs have shown protective effects in pancreatitis, though the precise mechanisms remain insufficiently explored (23).

Extracellular vehicles (EVs), which are released by various cell types, play critical roles in a range of biological processes(24). The therapeutic potential of EVs in pancreatic diseases has gained attention, and stem cell-derived EVs have been shown to alleviate AP symptoms(25). For example, hUC-MSC-derived EVs (hUC-MSC-EVs) inhibit necrosis in AP acinar cells(26), while EVs from hair follicle stem cells reduce inflammation and pyroptosis in these cells(27). Enhancing the therapeutic efficacy of stem cell-derived exosomes for AP treatment is a key area of research. For instance, treating hUC-MSCs with TNF-α has been shown to improve the efficacy of their EVs in alleviating AP(26).

Given that both emodin and hUC-MSC-EVs have demonstrated therapeutic potential in AP, we investigated whether emodin-loaded hUC-MSC-EVs could provide enhanced protection for inflamed pancreatic tissues. To explore this, we first purified hUC-MSC-EVs and generated emodin-loaded EVs. We then established AP cell, organoid, and mouse models using streptozotocin (STZ) to induce the disease. Finally, we compared the effects of emodin, hUC-MSC-EVs, and emodin-loaded hUC-MSC-EVs on cell viability, inflammation, and pyroptosis in these AP models.

## Materials and methods

### Cell culture

AR42J cells were obtained from IMMOCELL (Xiamen, China) and maintained in F-12K medium containing 20% fetal bovine serum and 1 x penicillin-streptomycin solution (Gibco, USA) in a humidified incubator at 37°C with 5% CO_2_.

### Characterization of hUC-MSCs

hUC-MSCs were obtained from IMMOCELL and cultured in MEM α medium (#12571-063, Thermo Fisher Scientific, USA). To confirm the mesenchymal stem cell (MSC) characteristics of these cells, they were incubated with fluorophore-conjugated antibodies targeting specific cell surface markers and then analyzed using a NovoCyte flow cytometer (Agilent, USA). The antibodies used for this analysis are as follows: FITC-conjugated anti-CD34 (#343503, 1:100), PE-conjugated anti-CD14 (#982508, 1:100), PE-conjugated anti-CD31 (#303105, 1:100), FITC-conjugated anti-CD105 (#800505, 1:100), PE-conjugated anti-CD45 (#304007, 1:100), PE-conjugated anti-CD73 (#344015, 1:100), PE-conjugated anti-CD90 (#328109, 1:100), PE-conjugated anti-CD29 (#303007, 1:100), FITC-conjugated anti-CD44 (#338803, 1:100), and a PE-conjugated isotype control antibody (ISO, #981804, 1:100). All antibodies were purchased from BioLegend (USA).

To further verify the pluripotency of the hUC-MSCs, we induced differentiation into chondrocytes, osteocytes, and adipocytes using the following kits: STEMPRO® Adipogenesis Differentiation Kit (#A1007001), STEMPRO® Chondrogenesis Differentiation Kit (#A1007101), and STEMPRO® Osteogenesis Differentiation Kit (#A1007201). All kits were purchased from Thermo Fisher Scientific, and the procedures strictly followed the manufacturer’s instructions.

### Alizarin red, Alcian blue, and Oil red staining

The hUC-MSCs underwent a 14-day induction process, followed by a rinse with phosphate-buffered saline (PBS). Next, the cells were fixed in 4% paraformaldehyde (PFA) for 30 minutes at room temperature (RT). For Alizarin red staining, the fixed cells were treated with 60% isopropanol for 1 minute at RT, followed by additional washes with PBS. The cells were then stained with a 10% Alizarin red solution for 15 minutes at RT, after which they were rinsed with PBS. For Alcian blue staining, the cells were incubated with a 1% Alcian blue solution in 0.1 N HCl for 30 minutes at RT, rinsed three times with 0.1 N HCl, and then placed in distilled water. For Oil Red O staining, the fixed cells were incubated with a 0.5% Oil Red O solution in isopropanol for 5 minutes at 37°C, protected from light. All staining results were observed and imaged using a BX51 inverted microscope equipped with a U-CMAD3 camera (OLYMPUS, Japan).

### EV purification and characterization

Conditioned media from hUC-MSCs underwent a series of centrifugation steps to isolate purified EVs. First, it was centrifuged at 2,000 g for 30 minutes at 4°C, followed by a second centrifugation of the supernatant at 10,000 g for another 30 minutes at 4℃. The supernatant was then collected and filtered through 0.45 μ m filters before undergoing further centrifugation at 100,000 g for 60 minutes at 4°C. The resulting pellet was resuspended in PBS, filtered through 0.22 μm filters, and centrifuged again at 100,000 g for 60 minutes at 4°C. Finally, the supernatant was carefully discarded, and the pellet was resuspended in 100 μL of ice-cold PBS.

For analysis, two 10 μL aliquots of EVs were set aside for transmission electron microscopy (TEM) and nanoparticle tracking analysis (NTA) to assess the morphology and size of the purified EVs. The remaining EV suspensions were stored at -80°C for future experiments. The presence of EV surface markers CD9, CD81, and CD63 was confirmed using Western blot analysis.

### TEM and NTA

TEM and NTA were performed by Pinuofei Biological Technology (China). For TEM sample preparation, the cell culture media were first removed, and the cells were washed with PBS before being fixed in 2.5% glutaraldehyde. Afterward, the fixed cells were digested, pelleted, and resuspended, followed by a second fixation with 2.5% glutaraldehyde for 2 hours at room temperature (RT).

### CCK-8 assay

Cell viability was assessed using the CCK-8 assay kit (C0038, Beyotime, China), following the manufacturer’s instructions. Briefly, cells were seeded at a density of 2,000 cells per well in a 96-well plate and incubated for 12 hours. Afterward, the cells were treated with 200 μM STC for the designated time period. Next, 10 μL of CCK-8 working solution was added to each well, and the cells were incubated in the dark for 1 hour. Finally, optical density (OD) readings at 450 nm were measured using a microplate reader.

### Carboxyfluorescein succinimidyl ester (CFSE) tracing

CFSE dye (40714ES76, Yeasen, China) was thoroughly mixed with the EVs at a 1:1000 ratio and incubated for 2 hours at 37°C. The mixture was then added to the cells and incubated for 48 hours. Afterward, the cells were washed with culture medium, counterstained with DAPI, mounted, and imaged using LSM 900 confocal microscopy (Zeiss, Germany).

### Western blot

Proteins were isolated using RIPA buffer supplemented with protease and phosphatase inhibitors (Beyotime), followed by homogenization and sonication on ice. Protein concentrations were determined using a BCA kit (TIANGEN Biotechnology, China). The proteins were then separated on 10% SDS-PAGE gels and transferred onto PVDF membranes. After blocking the membranes with 5% non-fat milk for 1 hour at room temperature (RT), they were incubated overnight at 4°C with primary antibodies. The next day, the membranes were rinsed with TBST and incubated with secondary antibodies for 1 hour at RT. Protein signals were visualized using an ECL kit (Beyotime) and detected with a Bio-Rad ChemiDoc imaging system. ImageJ software (NIH, USA) was used to quantify the intensity of the protein bands. Antibody details are provided in Table 1.

### Quantitative polymerase chain reaction (qPCR)

Total RNA was extracted using TRIzol reagent (Thermo Fisher Scientific). cDNA synthesis was performed using the HiScript II Q RT SuperMix for qPCR (R222-01, Vazyme, China). qPCR was conducted with ChamQ SYBR Color qPCR Master Mix (Q431-02, Vazyme) on a CFX Connect 96-well system (Bio-Rad, USA). Target gene expression levels were quantified using the 2−ΔΔCt method. The qPCR primer sequences are listed in Table 2.

### Measurement of pro-inflammatory cytokines and ATP

The concentrations of IL-1β, IL-18, and IL-6 in the culture medium or mouse serum were measured using the respective ELISA kits. ATP levels were also assessed using a specific kit. All measurements were performed according to the manufacturer’s instructions. The kits used for these analyses are listed in Table 3.

### Reactive oxygen species (ROS) level detection

ROS levels were measured using a DCF-DA kit (Yeason, 50101ES01, China), following the manufacturer’s instructions. Cells or organoids were digested, resuspended in serum-free medium, and incubated with 10 μM DCF-DA at 37°C for 20 minutes in the dark, with periodic inversion of the mixture. After incubation, the suspensions were rinsed with serum-free medium, pelleted to remove any unbound DCF-DA, resuspended in PBS, and analyzed by flow cytometry.

### Establishment of human pancreas organoid (HPO)

HPO was established following a modified protocol from a previous study using the human embryonic stem cell (hESC) line H1(28). Two essential culture media, BE1 and BE2, were used to induce HPO. BE1 consists of 0.1% BSA, 2 mM L-glutamine, and DMEM/F12. BE2 contains 2% BSA, 2 mM L-glutamine, 44 mg/mL ascorbic acid, 0.5x Insulin-Transferrin-Selenium-Ethanolamine, and DMEM/F12.

To induce pancreatic differentiation, cells were cultured in BE1 medium containing 3 mM CHIR99021 and 100 ng/mL Activin for one day (day 0). For the next three days (day 1–3), the medium was switched to BE1 + 100 ng/mL Activin, and refreshed daily to promote definitive endoderm (DE) formation. On days 4 and 5, the culture medium was changed to BE1 + 50 ng/mL KGF to induce gut tube endoderm (GTE). From days 6 to 9, the medium was switched to BE2 + 0.25 mM SANT-1, 2 mM retinoic acid, 200 nM LDN-193189, and 500 nM PD0325901 to complete pancreatic endoderm (PE) differentiation.

From days 10 to 13, the medium was changed to BE2 containing 50 ng/mL FGF10, 330 nM Indolactam V, 10 mM SB431542, and 16 mM glucose to induce pancreatic progenitor (PP) differentiation. On day 12, the cells were digested with TrypLE enzyme for 5 minutes at 37°C, then resuspended in DMEM/F12 medium, centrifuged, and resuspended in full culture medium containing 10 μM Y27632. The suspension was mixed with growth factor-reduced Matrigel (Corning, USA) at a 1:3 ratio on ice, and 30 μL of the mixture was plated into 24-well plates and incubated at 37°C for 20 minutes. Afterward, 500 μL of Y27632-containing medium was added to each well. On day 14, cells were cultured in BE2 + 10 ng/mL FGF2 + 10 mM Y27632 until day 17. From day 18 onward, the organoids were cultured in BE2 + 10 ng/mL FGF2 + 10 mM nicotinamide, with the medium refreshed every 2 to 3 days.

### Establishment of AP mouse model

Eight-week-old male CD-1 mice, weighing between 25 g and 28 g, were obtained from Beijing Vital River Laboratory Animal Technology Co., Ltd. The mice were randomly divided into five groups: Sham, AP, AP+emodin, AP+hUC-MSC-EV, and AP+hUC-MSC-EV+emodin, with 10 mice per group. The mice underwent one week of acclimation in a facility with a 12-hour light/12-hour dark cycle. All mice were fasted for 12 hours prior to AP surgery. For the groups receiving emodin or EV treatment, mice were injected via the tail vein with either emodin (25 mg/kg body weight), hUC-MSC-EVs (300 μg/mL, 100 μL/mouse), or hUC-MSC-EVs encapsulated with emodin (100 μL/mouse) 30 minutes before surgery. Emodin-loaded EVs were prepared by mixing hUC-MSC-EVs (600 μg/mL) with emodin (5 mg/mL) at a 1:1 ratio on a vortex mixer at 37°C for 60 minutes.

During surgery, the mice were anesthetized with 3% isoflurane inhalation and positioned on a heating pad. A midline laparotomy was performed to access the abdominal cavity, and the bile-pancreatic (BP) duct was occluded with a vascular clamp at the hepatic hilum. The BP duct near the duodenal end was ligated 2 mm from the needle injection site. A 31-gauge needle was then used to inject either 2 mL/kg of 5% sodium taurocholate into the BP duct for the AP groups, or saline for the Sham group.

Twenty-four hours post-surgery, the mice were euthanized, and blood and pancreatic tissue samples were collected. Serum samples were used for ELISA assays, and pancreatic tissue was either fixed in 4% PFA for histological analysis or frozen at -80° C for subsequent qPCR and Western blot assays. All experimental procedures were approved by the Institutional Ethics Committee of the First Affiliated Hospital of Xiamen University (XMULAC20240193).

### Immunohistochemistry (IHC) and immunofluorescence (IF)

For immunohistochemistry (IHC), organoids were fixed in 4% PFA and sectioned into thick slices. These slices were dewaxed, rehydrated, and subjected to antigen retrieval, followed by treatment with 3% H2O2. The sections were then permeabilized with 0.2% Triton X-100 in PBS and blocked with 1% BSA. Afterward, the sections were incubated with primary antibodies, washed with PBST, and then incubated with secondary antibodies for 1 hour at room temperature (RT). Following another wash, the sections were stained with DAB working solution (Beyotime), counterstained with hematoxylin, and mounted. Images were captured using a BX51 inverted microscope, and the average optical density (AOD) was quantified using Image-Pro Plus 6.0.

For immunofluorescence (IF), fixed HPOs were embedded in OCT and sectioned into 10 μm thick slices using a cryostat. The slides were rinsed with PBST and blocked with 1% BSA for 1 hour. They were then incubated with primary antibodies overnight at 4°C, followed by three washes with PBST. Next, the slides were incubated with secondary antibodies for 1 hour at RT, followed by three more washes with PBST. Finally, the slides were counterstained with DAPI, mounted, and observed using LSM 900 confocal microscopy (Zeiss, Germany).

The following antibodies were used for IHC or IF: anti-AMY (66133-1-lg, Proteintech, 1:100), anti-CK19 (14965-1-AP, Proteintech, 1:100), anti-ECAD (20874-1-AP, Proteintech, 1:100), anti-Ki-67 (A21861, ABclonal, 1:200), anti-Caspase-1 (abs119750, Absin, 1:200), anti-ASC (10500-1-AP, Proteintech, 1:200), HRP Goat Anti-Mouse/Rabbit IgG (RS0011, Immunoway, ready to use), and Alexa Fluor 488-conjugated goat anti-rabbit IgG (RS3211, Immunoway, 1:500).

### Statistical analysis

Data analysis and visualization were performed using GraphPad Prism software (v8.0, USA). Results are presented as mean ± SD. An unpaired two-tailed Student’s t-test was used to assess significance between two groups. For comparisons among three or more groups, statistical significance was determined using one-way analysis of variance (ANOVA), with a p-value of less than 0.05 considered statistically significant.

## Results

### Characterization of hUC-MSCs and hUC-MSC-EVs

To validate the pluripotency of the hUC-MSCs, we first assessed their identity through flow cytometry. The results showed that these cells expressed MSC surface markers CD29, CD105, CD44, CD73, and CD90, while being negative for CD34, CD31, CD45, and CD14 (Figure S1A). Next, we induced differentiation into osteocytes, chondrocytes, and adipocytes using specific culture media. Successful differentiation was confirmed by Alizarin Red, Alcian Blue, and Oil Red staining for osteocytes, chondrocytes, and adipocytes, respectively (Figure S1B). These findings demonstrate the pluripotent characteristics of the hUC-MSCs.

To isolate EVs from these hUC-MSCs, we conducted multiple centrifugation steps and analyzed the purified EVs. The results showed that the EVs exhibited typical morphology, with a size distribution ranging from 20 to 120 nm (Figure S2A and S2B). Consistent with these observations, Western blot analysis confirmed the presence of EV markers CD9, CD81, and CD63 (Figure S2C and S2D). Likewise, emodin-loaded hUC-MSC-EVs also exhibited typical EV morphology, as evidenced by TEM (Figure S2E). Collectively, these findings confirm the successful purification of hUC-MSC-EVs.

### Emodin Enhances the Effects of hUC-MSC-EVs in Promoting Viability, Reducing Inflammation, and Inhibiting Pyroptosis in STZ-Treated AR42J Cells

To investigate the impact of hUC-MSC-EVs on inflammatory pancreatic cells, we first used AR42J cells, which are epithelial-like cells derived from a rat pancreatic tumor(29). By tracking CSFE-labeled hUC-MSC-EVs, we found that these cells efficiently absorbed the purified EVs (Figure S2F).

We then treated the AR42J cells with STZ, a chemical that damages pancreatic cells and induces diabetes(30), over various time periods. Using MTT assays to measure cell viability, we observed that AR42J cells exposed to 200 μM STZ for 12 hours showed a significant reduction in viability (Figure S3A). Moreover, these cells secreted significantly higher levels of TNF-α, IL-1β, and IL-18 (Figure S3B). Based on these findings, we decided to treat AR42J cells with 200 μM STZ for 12 hours to establish a cell model of AP. We also optimized the dose of emodin and found that 5 mg/mL was sufficient to reduce cell viability (Figure S3C). As a result, we selected 2.5 mg/mL as the emodin concentration for subsequent analyses.

Next, we cultured the AP cell model with emodin, hUC-MSC-EVs, or emodin-loaded hUC-MSC-EVs (hUC-MSC-EV+emodin). Compared to AP cells treated with PBS, emodin, hUC-MSC-EVs, and hUC-MSC-EV+emodin all significantly improved the viability of STZ-treated AR42J cells, with hUC-MSC-EV+emodin demonstrating a more pronounced pro-viability effect than hUC-MSC-EVs alone (Figure 1A). Similarly, hUC-MSC-EV+emodin outperformed hUC-MSC-EVs in reducing ROS levels in STZ-treated AR42J cells, as indicated by DCF-DA staining (Figure 1B and C). Furthermore, emodin-loaded hUC-MSC-EVs exhibited a stronger anti-inflammatory effect than control hUC-MSC-EVs in these AP cells, as demonstrated by the mRNA expression and secretion levels of TNF-α, IL-1β, and IL-18 measured by qPCR and ELISA (Figure 1D and E). Additionally, the mitochondrial structure was more intact in AP cells treated with hUC-MSC-EV+emodin, although hUC-MSC-EVs alone also effectively repaired STZ-induced mitochondrial damage (Figure 1F).

**Figure 1.**
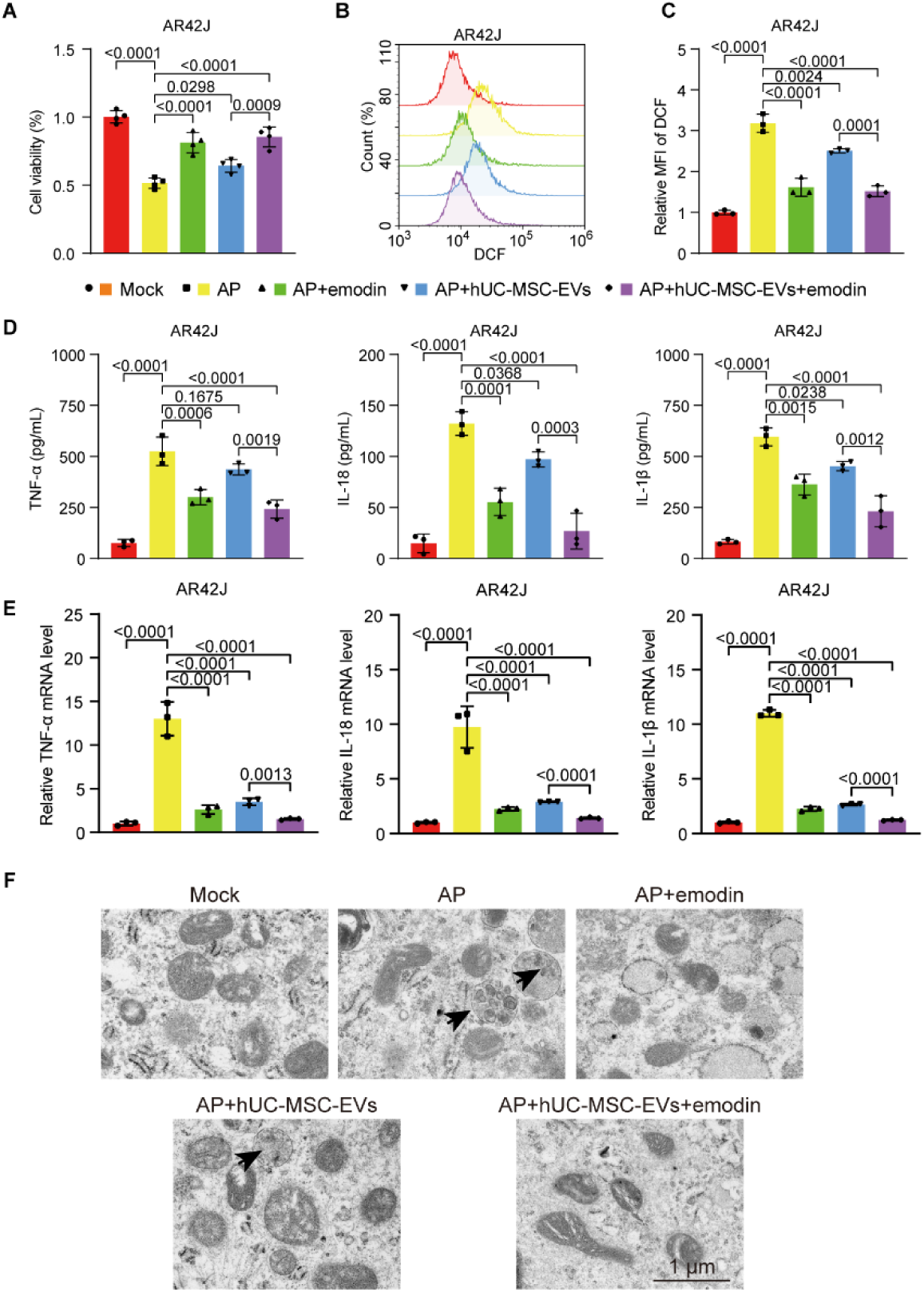
Emodin strengthens the effect of hUC-MSC-EVs in promoting viability, reducing pro-inflammatory cytokine secretion, and decreasing ROS levels in AR42J cells exposed to STZ. (A) MTT assay data showing the viability of STZ-induced AR42J cells after the indicated treatments. “Mock” refers to cells treated with an equivalent amount of PBS used for STZ dilution. (B) DCF-DA staining results depicting ROS levels in AR42J cells after the indicated treatments. (C) Quantification of DCF-DA staining results. (D) ELISA data indicating TNF-α, IL-1β, and IL-18 levels in the conditioned media after the indicated treatments. (E) qPCR results illustrating mRNA levels of TNF-α, IL-1β, and IL-18 in AR42J cells after the indicated treatments. (F) Representative TEM images showing mitochondrial morphology in AR42J cells after the indicated treatments. Black arrows indicate damaged mitochondria.

Given the close association between inflammation and pyroptosis, we then evaluated pyroptosis markers in these cells via Western blot. Our data showed that emodin, hUC-MSC-EVs, and emodin-loaded hUC-MSC-EVs significantly reduced the levels of NLRP3, ASC, Cleaved-Caspase 1, and Cleaved-GSDMD, with emodin-loaded hUC-MSC-EVs demonstrating the greatest efficacy (Figure 2A and B). These findings suggest that emodin encapsulation enhances the anti-inflammatory and anti-pyroptotic functions of hUC-MSC-EVs in STZ-treated AR42J cells.

**Figure 2.**
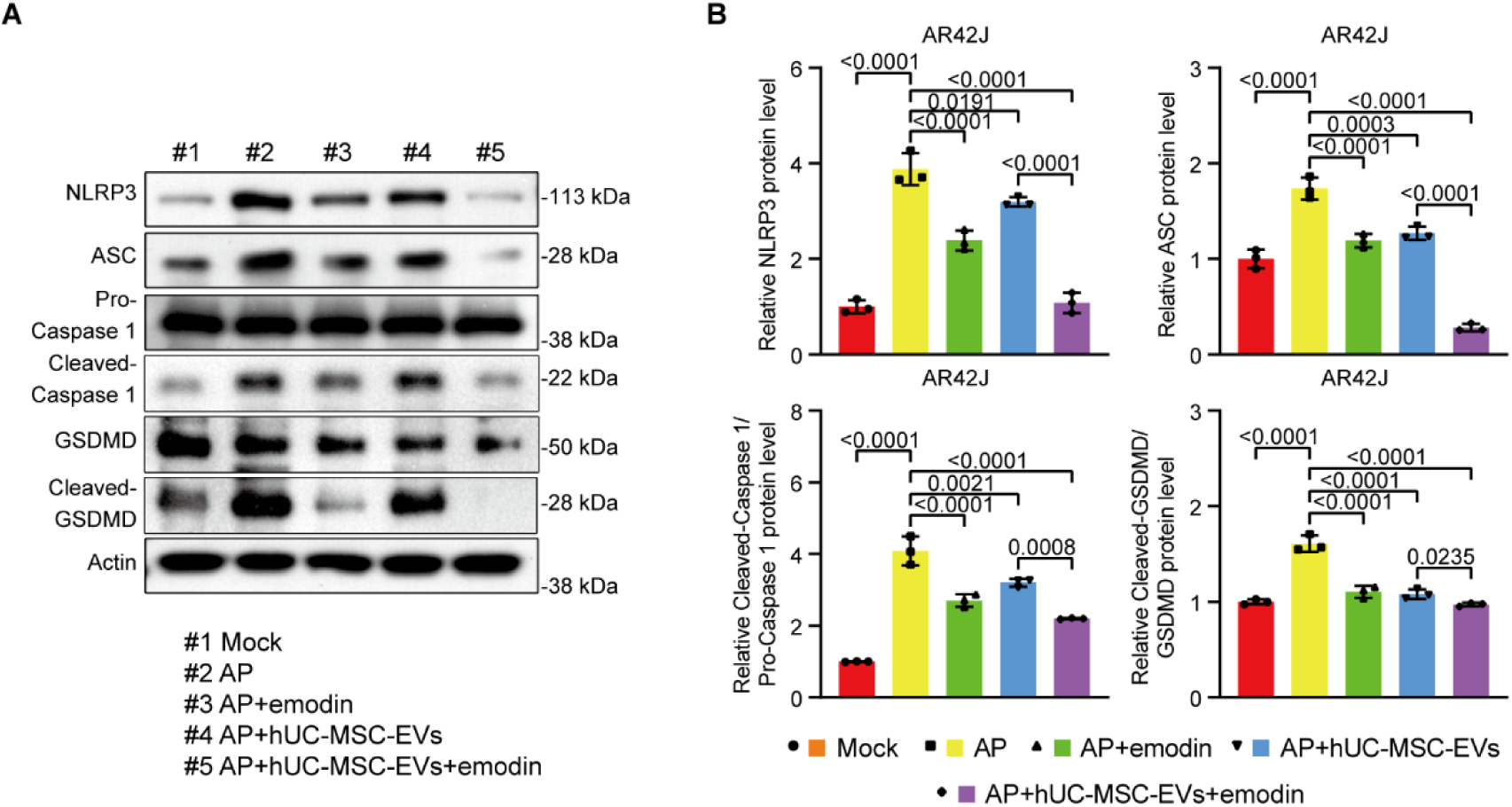
Emodin enhances the anti-pyroptotic function of hUC-MSC-EVs in STZ-treated AR42J cells. (A) Western blot data showing the expression of NLRP3, ASC, Pro-Caspase 1, Cleaved-Caspase 1, GSDMD, and Cleaved-GSDMD in AR42J cells after the indicated treatments. (B) Quantification of Western blot data.

### Emodin Enhances the Protective Effects of hUC-MSC-EVs in STZ-Treated HPOs”

Next, we validated our findings from AR42J cells using an HPO model. Initially, we established HPOs from hESCs, which exhibited significant morphological changes during differentiation (Figure S4A). Both IF and qPCR assays confirmed that the differentiating HPOs expressed pancreatic exocrine markers AMY, AMY2A, and CTRC, ductal markers CK19, SOX9, and CFTR, the epithelial marker ECAD, and progenitor markers PDX1 and NKX6.1 (Figure S4B and C)(28).

After establishing the HPOs, we treated them with STZ and various doses of emodin for 48 hours and assessed ATP levels, which reflect organoid viability(31). Our data showed that 10 mg/L of emodin exhibited significant cytotoxicity in HPOs (Figure S3D). Therefore, we used 5 mg/L emodin in subsequent experiments.

Similar to STZ-treated AR42J cells, emodin, hUC-MSC-EVs, and emodin-loaded hUC-MSC-EVs all significantly increased ATP levels in HPOs, indicating their pro-viability effects (Figure 3A). However, the ATP levels in HPOs treated with hUC-MSC-EV+emodin were higher than those treated with control hUC-MSC-EVs (Figure 3A). Consistently, emodin-loaded hUC-MSC-EVs were more effective than control EVs in reducing ROS levels in STZ-treated HPOs (Figure 3B and C). Furthermore, HPOs cultured with emodin-loaded hUC-MSC-EVs exhibited lower expression and secretion of TNF-α, IL-1β, and IL-18 compared to those treated with control hUC-MSC-EVs (Figure 3D and E). Additionally, the increased levels of NLRP3, ASC, Cleaved-Caspase 1, and Cleaved-GSDMD induced by STZ in HPOs were reduced by treatments with emodin, hUC-MSC-EVs, or emodin-loaded hUC-MSC-EVs, with the latter showing the greatest efficacy (Figure 4A and B). These results demonstrate that emodin enhances the pro-viability, anti-inflammatory, and anti-pyroptotic functions of hUC-MSC-EVs in STZ-treated HPOs.

**Figure 3.**
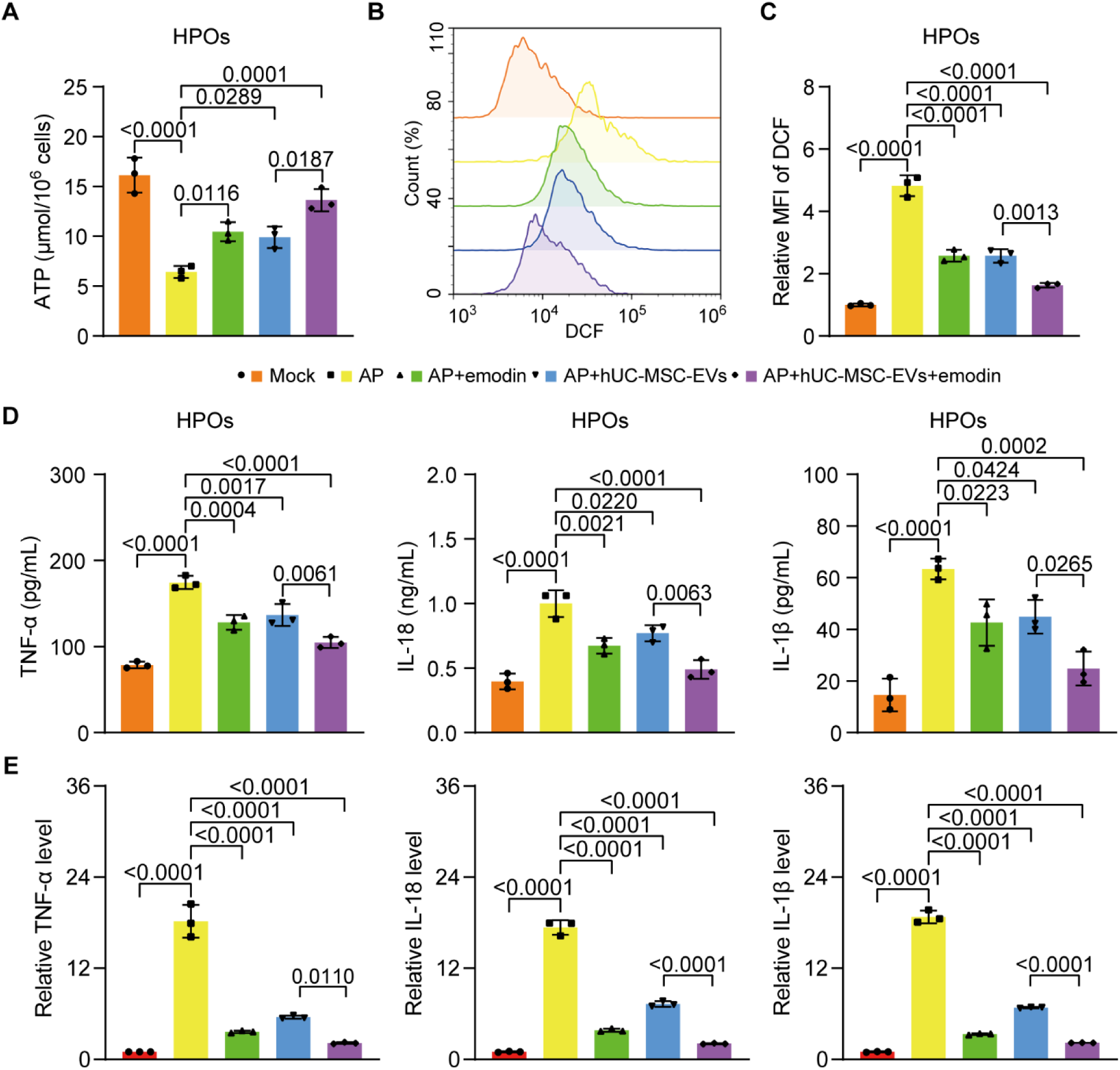
Emodin-loaded hUC-MSC-EVs exhibit the best efficacy in promoting viability, reducing ROS levels, and decreasing pro-inflammatory cytokine secretion in STZ-treated HPOs. (A) ATP levels in STZ-induced HPOs after the indicated treatments. (B) DCF-DA staining results showing ROS levels in HPOs after the indicated treatments. (C) Quantification of DCF-DA staining results. (D) ELISA data indicating TNF-α, IL-1β, and IL-18 levels in the conditioned media of HPOs after the indicated treatments. (E) qPCR results illustrating mRNA levels of TNF-α, IL-1β, and IL-18 in HPOs after the indicated treatments.

**Figure 4.**
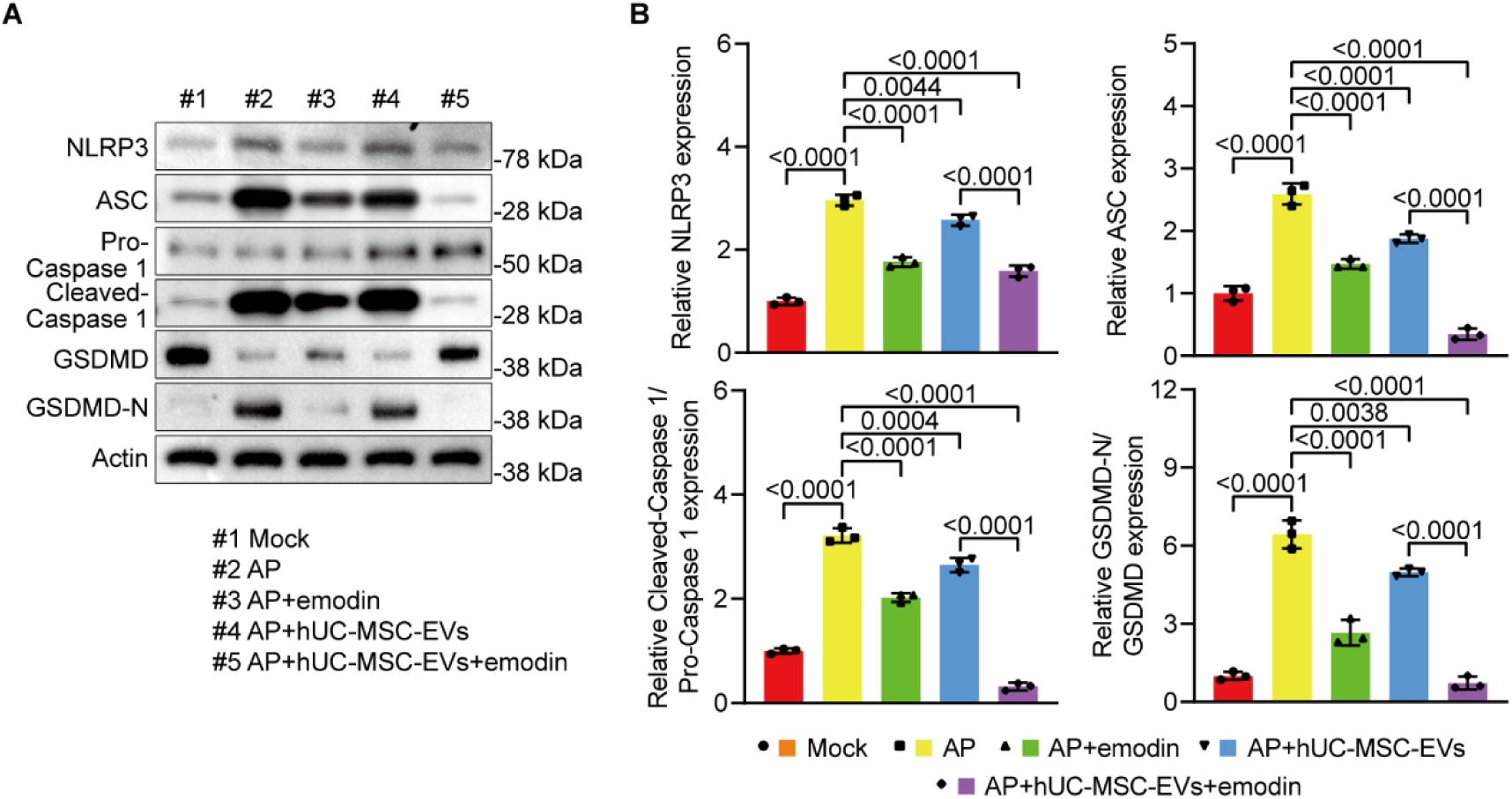
Emodin bolsters the anti-pyroptotic function of hUC-MSC-EVs in STZ-treated HPOs. (A) Western blot data showing the expression of NLRP3, ASC, Pro-Caspase 1, Cleaved-Caspase 1, GSDMD, and Cleaved-GSDMD in AR42J cells after the indicated treatments. (B) Quantification of Western blot data.

### Emodin Enhances the Effects of hUC-MSC-EVs in an AP Mouse Model

To further investigate the protective effects of emodin and hUC-MSC-EVs on inflamed pancreatic tissues, we used an AP mouse model. H&E staining confirmed that treatments with emodin, hUC-MSC-EVs, or emodin-loaded hUC-MSC-EVs significantly alleviated pancreatic tissue damage in the AP mice (Figure 5A). Consistently, IHC data showed that the reduced proliferation, indicated by Ki-67 expression, was improved by all treatments. Notably, pancreatic tissues from AP mice treated with emodin-loaded hUC-MSC-EVs exhibited higher proliferation compared to those treated with control hUC-MSC-EVs (Figure 5B and C). Additionally, serum levels and pancreatic mRNA expression of TNF-α, IL-1β, and IL-18 were significantly downregulated by all treatments, with emodin-loaded hUC-MSC-EVs demonstrating the greatest reduction in these pro-inflammatory cytokines (Figure 5D and E). At the molecular level, hUC-MSC-EVs loaded with emodin also showed the highest efficacy in reducing pyroptosis markers, including ASC, GSDMD, Caspase 1, and NLRP3, as indicated by Western blot and IF results (Figure 6A-D). These findings highlight that emodin significantly enhances the protective effects of hUC-MSC-EVs on inflamed pancreatic tissues in vivo.

**Figure 5.**
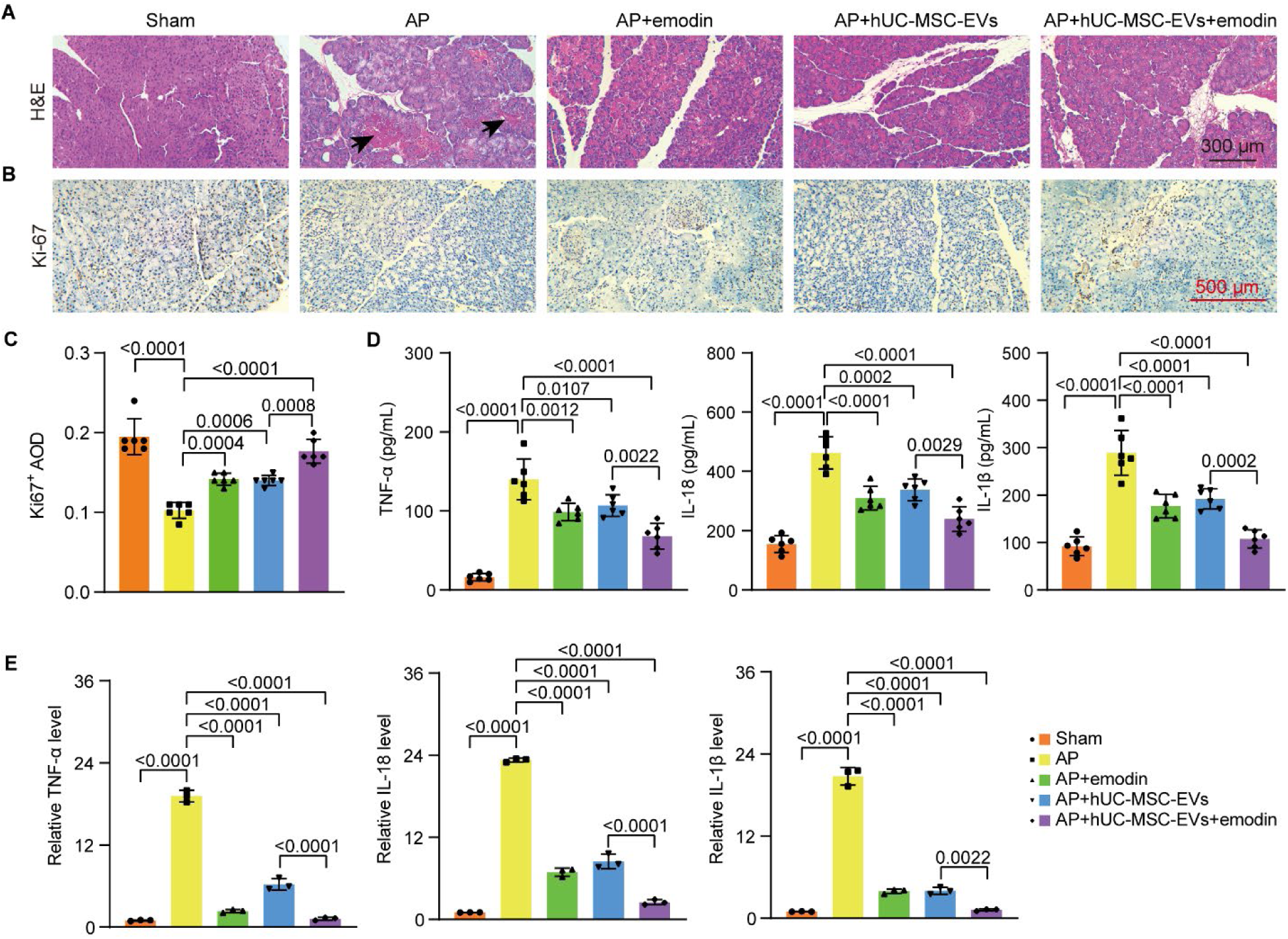
Emodin improves the effect of hUC-MSC-EVs in rescuing pancreatic cell proliferation and reducing pro-inflammatory cytokine secretion in AP mice. (A) H&E images showing the histology of pancreatic tissues from AP mice after the indicated treatments. Black arrows indicate damaged tissues. (B) IHC data depicting Ki-67 expression in pancreatic tissues from AP mice after the indicated treatments. (C) Quantification of Ki-67 IHC data. (D) ELISA data indicating serum levels of TNF-α, IL-1β, and IL-18 in AP mice after the indicated treatments. (E) qPCR results illustrating mRNA levels of TNF-α, IL-1β, and IL-18 in pancreatic tissues from AP mice after the indicated treatments.

**Figure 6.**
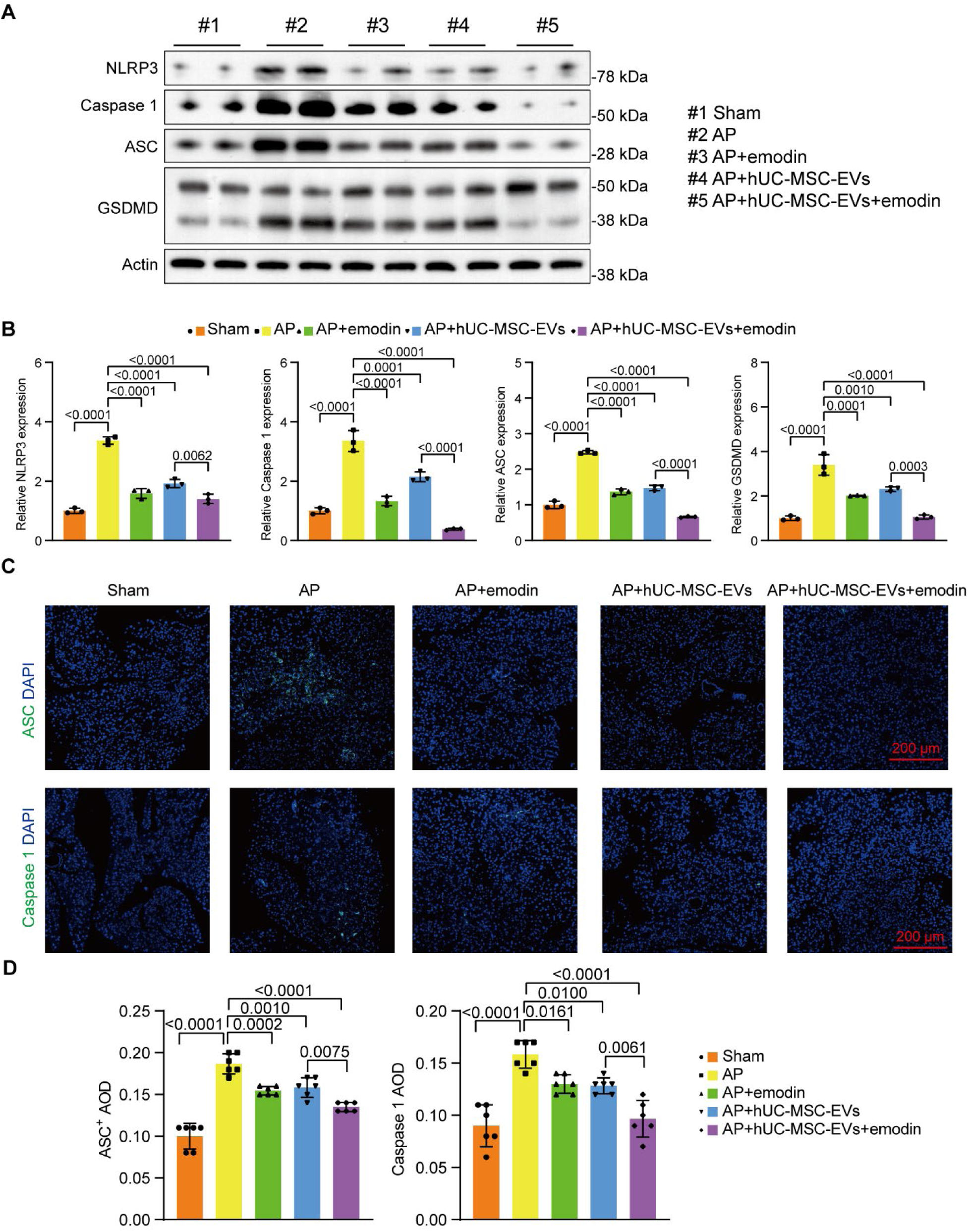
Emodin enhances the anti-pyroptotic effect of hUC-MSC-EVs in AP mice. (A) Western blot data showing the expression of NLRP3, Caspase 1, ASC, and GSDMD in pancreatic tissues from AP mice. (B) Results of Western blot data analysis. (C) IF results depicting ASC and Caspase 1 expression in pancreatic tissues from AP mice. (D) Quantification of IF results.

## Discussion

Our study introduces a novel and highly innovative approach to treating acute pancreatitis (AP) by leveraging the synergistic potential of emodin-loaded human umbilical cord mesenchymal stem cell-derived extracellular vesicles (hUC-MSC-EVs). This work not only reinforces the therapeutic efficacy of stem cell-derived EVs but also pioneers the engineering of these vesicles to enhance their protective effects by incorporating bioactive compounds like emodin. The combination of emodin and hUC-MSC-EVs represents a breakthrough strategy that directly addresses the limitations of current AP treatments(2), which primarily focus on supportive care without targeting the underlying inflammatory and pyroptotic mechanisms that drive disease progression.

The primary innovation of this study lies in the enhanced efficacy of hUC-MSC-EVs when loaded with emodin, a well-characterized anti-inflammatory compound derived from traditional Chinese medicine. Our data show that this combination significantly amplifies the protective effects of hUC-MSC-EVs, particularly in reducing inflammation, inhibiting pyroptosis, and improving cell viability in various AP models, including AR42J cells, human pancreatic organoids (HPOs), and an AP mouse model. These findings provide a promising alternative to conventional stem cell therapies, which are often limited by ethical concerns, immunogenicity, and tumorigenicity(32).

One of the most compelling aspects of our research is the demonstrated ability of emodin-loaded hUC-MSC-EVs to more effectively mitigate inflammatory responses compared to either treatment alone. The substantial reduction in pro-inflammatory cytokines, such as TNF-α, IL-1 β, and IL-18, alongside the decreased levels of pyroptosis markers (NLRP3, ASC, Cleaved-Caspase 1, and Cleaved-GSDMD), underscores the targeted action of this combination therapy. By specifically addressing the molecular pathways implicated in AP pathogenesis, this approach offers a significant advancement over existing therapies, which are largely nonspecific and fail to prevent disease progression at the cellular level.

Additionally, our focus on pyroptosis, a form of programmed cell death distinct from apoptosis and necrosis, marks a novel contribution to AP research. While pyroptosis has been increasingly recognized as a key contributor to the inflammatory damage observed in AP, few studies have explored therapeutic interventions that specifically target this pathway(33-35). Our findings show that emodin-loaded hUC-MSC-EVs significantly reduce pyroptosis markers, indicating not only symptomatic relief but also a deeper molecular intervention aimed at halting the progression of AP. This mechanistic insight positions our study as a pioneering effort in targeting pyroptosis within the broader context of pancreatic inflammation.

The clinical implications of these findings are substantial. By demonstrating that emodin-loaded hUC-MSC-EVs can enhance cell viability, reduce reactive oxygen species (ROS), and suppress the secretion of inflammatory cytokines, our study lays the groundwork for the development of cell-free, targeted therapies for AP. This approach has the added advantage of circumventing many of the risks associated with traditional stem cell therapies, such as immune rejection and tumor formation(36). Moreover, the ability to engineer EVs with specific bioactive compounds opens up new possibilities for personalized medicine, where therapies could be tailored to target specific inflammatory pathways or disease mechanisms in individual patients.

While our research makes significant strides in advancing the therapeutic potential of hUC-MSC-EVs and emodin, several questions remain. Although we have clearly demonstrated the protective effects of this combination on inflammation and pyroptosis, it is still unclear whether hUC-MSC-EVs and emodin can also protect against other forms of cell death, such as apoptosis and ferroptosis. These processes, particularly apoptosis, are known to contribute to pancreatic tissue damage in AP, often driven by excessive ROS levels(37, 38). Future studies should explore whether the observed reduction in ROS levels by emodin-loaded EVs can extend to other cell death pathways, providing a more comprehensive protective effect against AP-induced damage.

Moreover, the pancreas is composed of diverse cell types, including acinar, ductal, and islet cells, each of which plays a unique role in pancreatic function and may respond differently to inflammatory stimuli. While our study has focused primarily on acinar cells, understanding how hUC-MSC-EVs and emodin affect other pancreatic cell types will be critical for optimizing this therapy and maximizing its therapeutic efficacy. Further research is needed to clarify the specific interactions between these treatments and different cell populations within the pancreas, as this will provide deeper insights into how emodin-loaded EVs might protect or regenerate damaged pancreatic tissue.

In conclusion, our study presents a groundbreaking therapeutic strategy for treating AP by combining the synergistic effects of emodin and hUC-MSC-EVs. This combination represents a major advancement in both the use of EVs as therapeutic agents and the integration of traditional medicine compounds into modern biomedical approaches. By effectively targeting inflammation and pyroptosis (Figure 7), this strategy offers a novel, highly effective treatment option for AP. Moreover, our findings pave the way for future innovations in the use of engineered EVs for treating other inflammatory and degenerative diseases. Continued investigation into the underlying molecular mechanisms and broader applications of this approach will be essential for translating these promising results into clinical practice.

**Figure 7.**
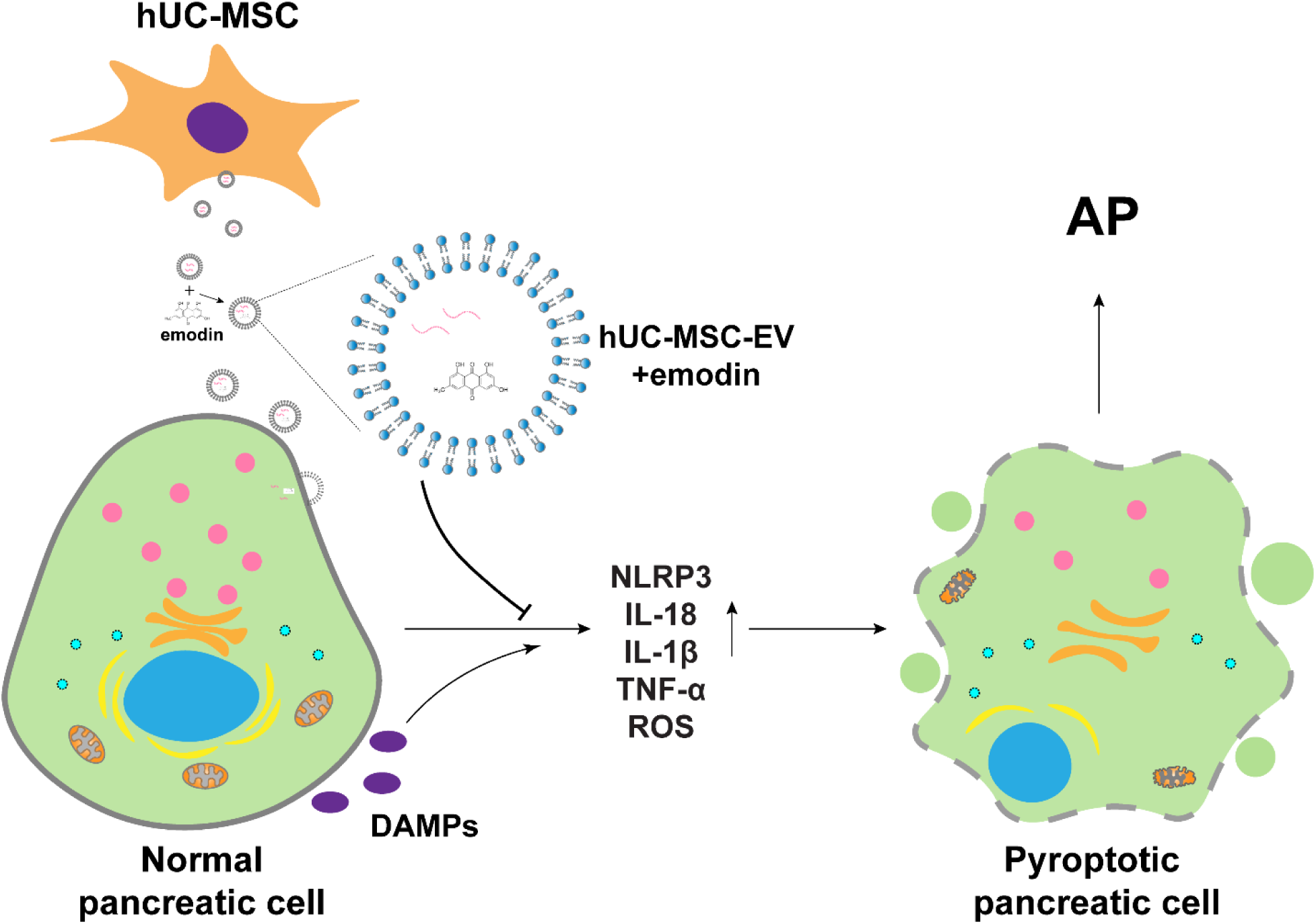
Schematic presentation of the protective role of hUC-MSC-EVs and emodin in pancreatic cells.

## Supplemental Figures

**Figure S1.**
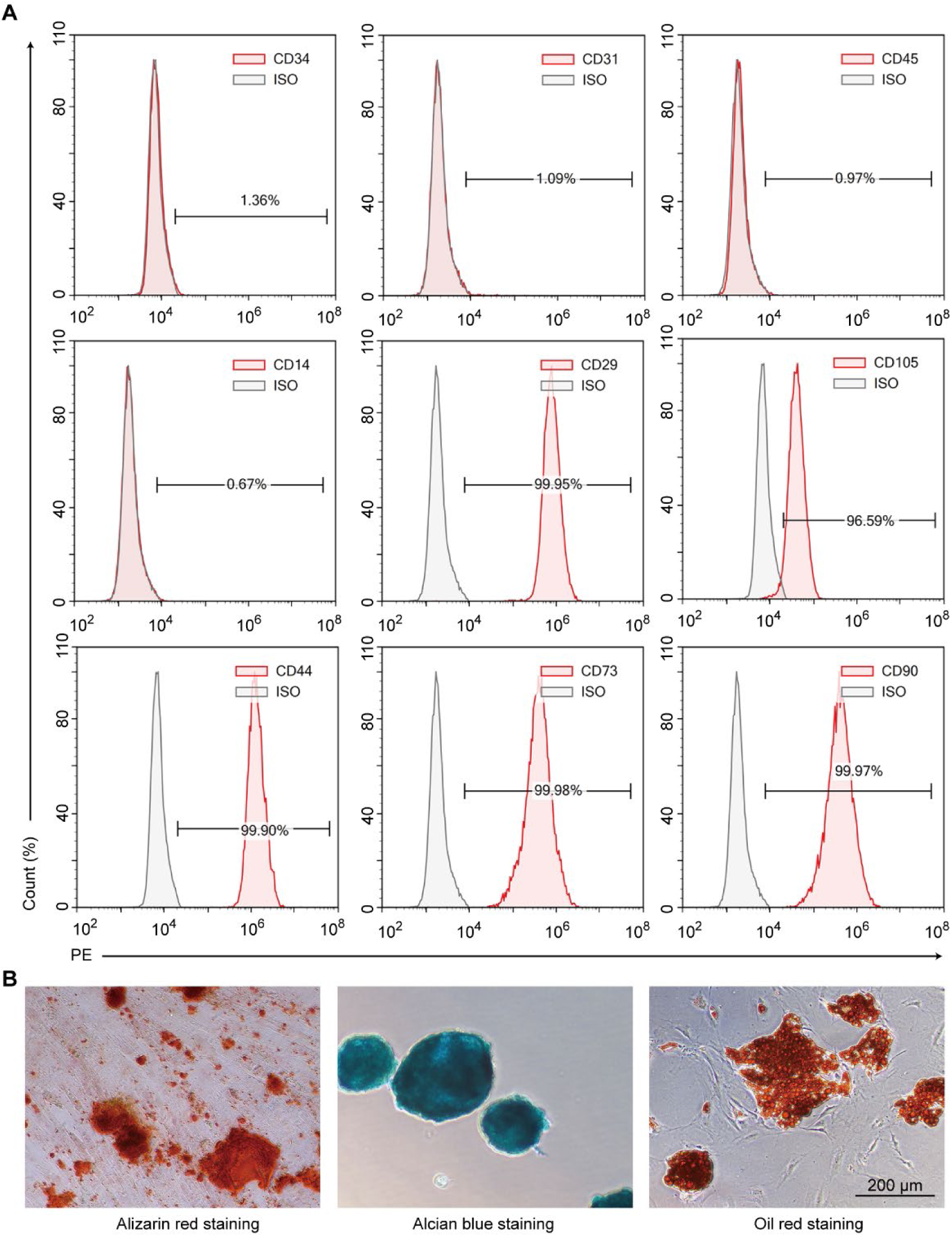
Characterization of hUC-MSCs. (A) Flow cytometry data showing negative expression of hematopoietic cell surface markers CD34, CD31, CD45, and CD14, and positive expression of MSC surface markers CD29, CD105, CD44, CD73, and CD90 on hUC-MSCs. (B) The effective differentiation of hUC-MSCs into osteocytes, chondrocytes, and adipocytes is confirmed by the positive staining results of Alizarin red, Alcian blue, and Oil red, respectively.

**Figure S2.**
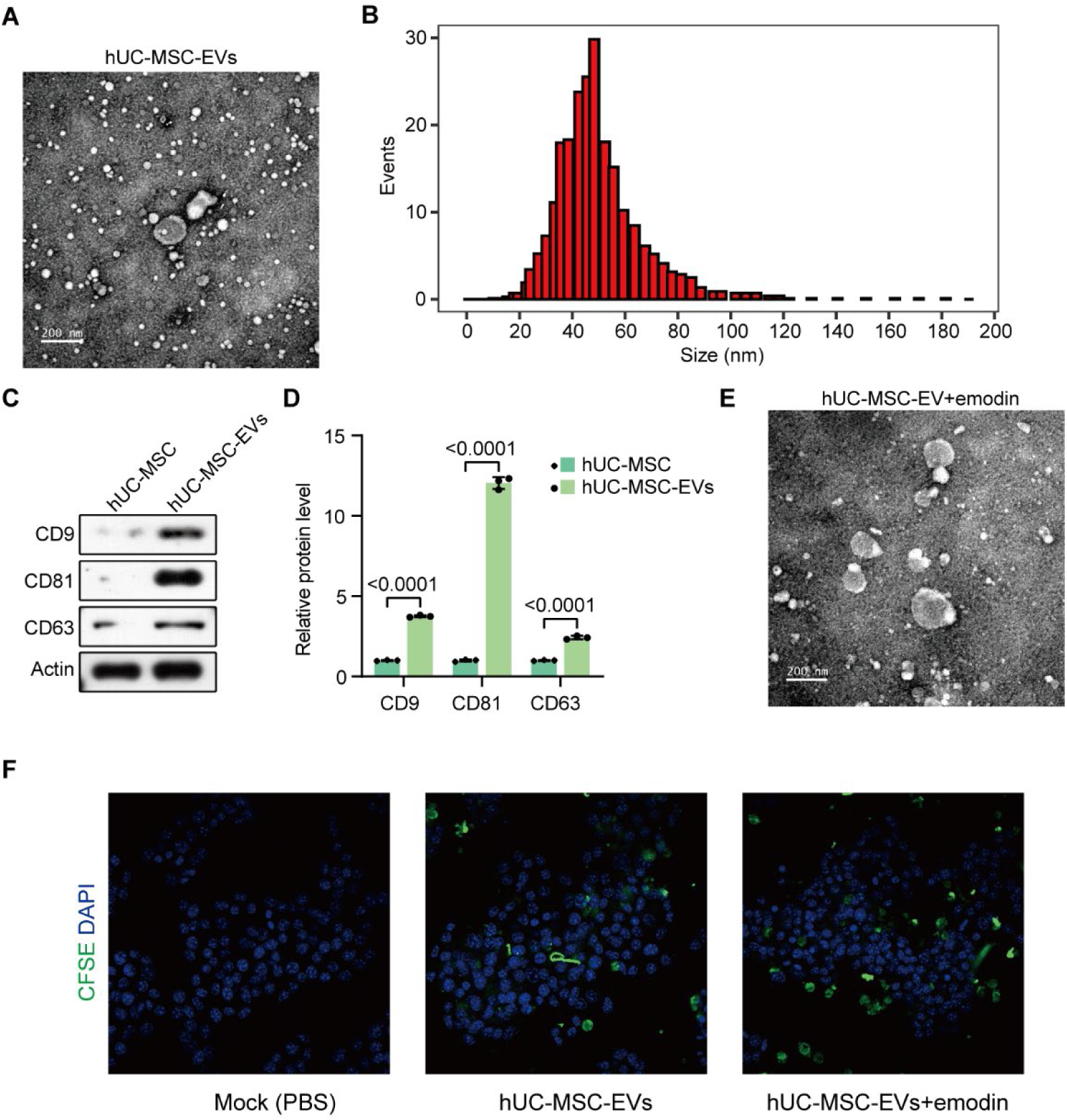
Identification of purified hUC-MSC-EVs. (A) TEM image showing the morphology of purified hUC-MSC-EVs. (B) NTA results depicting the size distribution of purified hUC-MSC-EVs. (C) Western blot data illustrating enrichment of CD9, CD81, and CD63 in purified hUC-MSC-EVs compared to hUC-MSCs. (D) Results of Western blot data analysis. (E) TEM image showing the morphology of emodin-loaded hUC-MSC-EVs. (F) Confocal images show the absorption of CFSE-labeled hUC-MSC-EVs and emodin-loaded hUC-MSC-EVs by AR42J cells.

**Figure S3.**
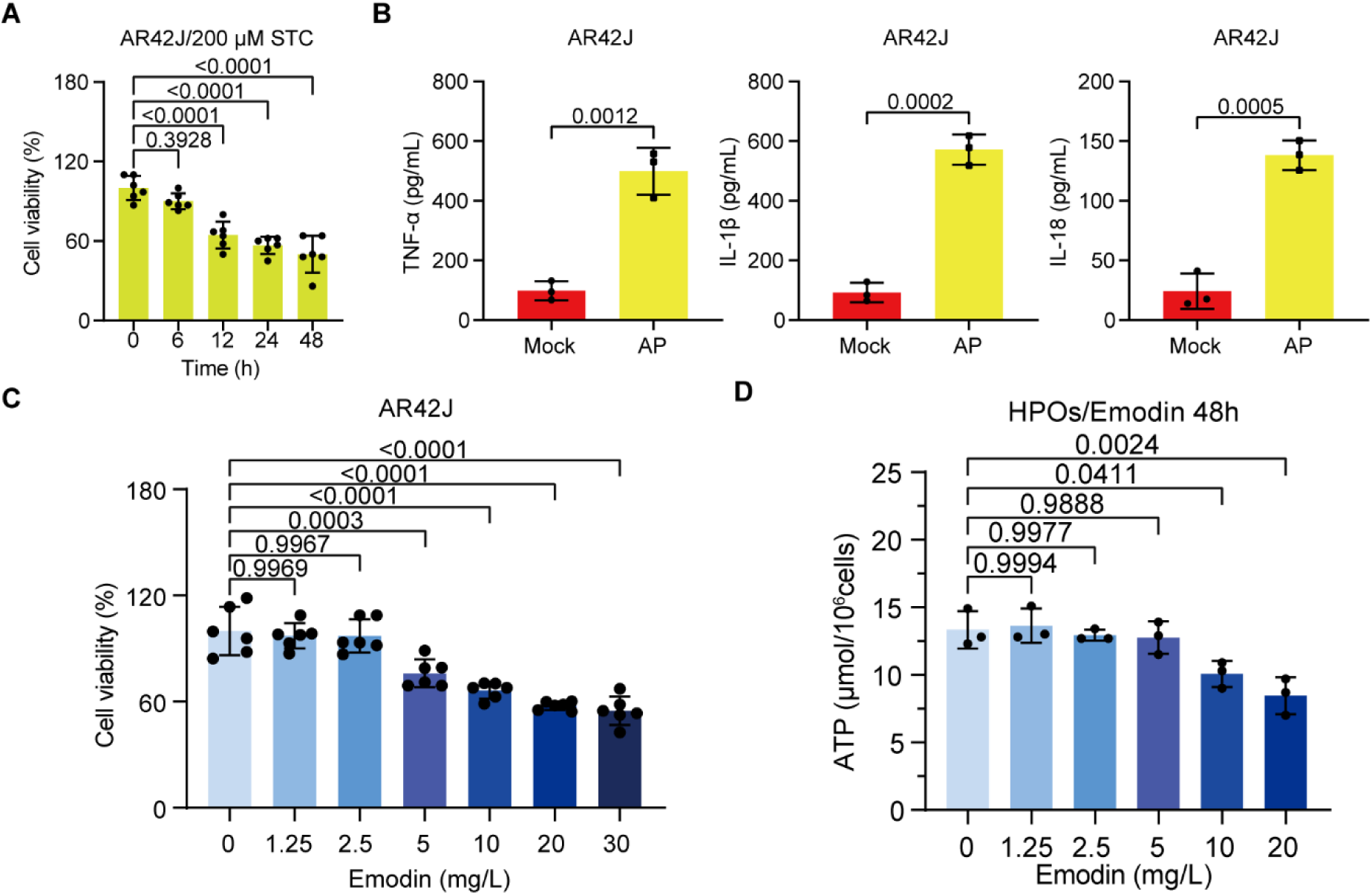
Optimization of STZ and emodin treatments. (A) MTT assay data showing the viability of cells exposed to 200 μM STZ for the indicated durations. (B) ELISA data indicating TNF-α, IL-1β, and IL-18 levels in the conditioned media of AR42J cells exposed to 200 μM STZ for 12 hours. (C) MTT assay results showing the viability of AR42J cells after treatment with specified concentrations of emodin. (D) ATP levels in HPOs following 48 hours of treatment with specified doses of emodin.

**Figure S4.**
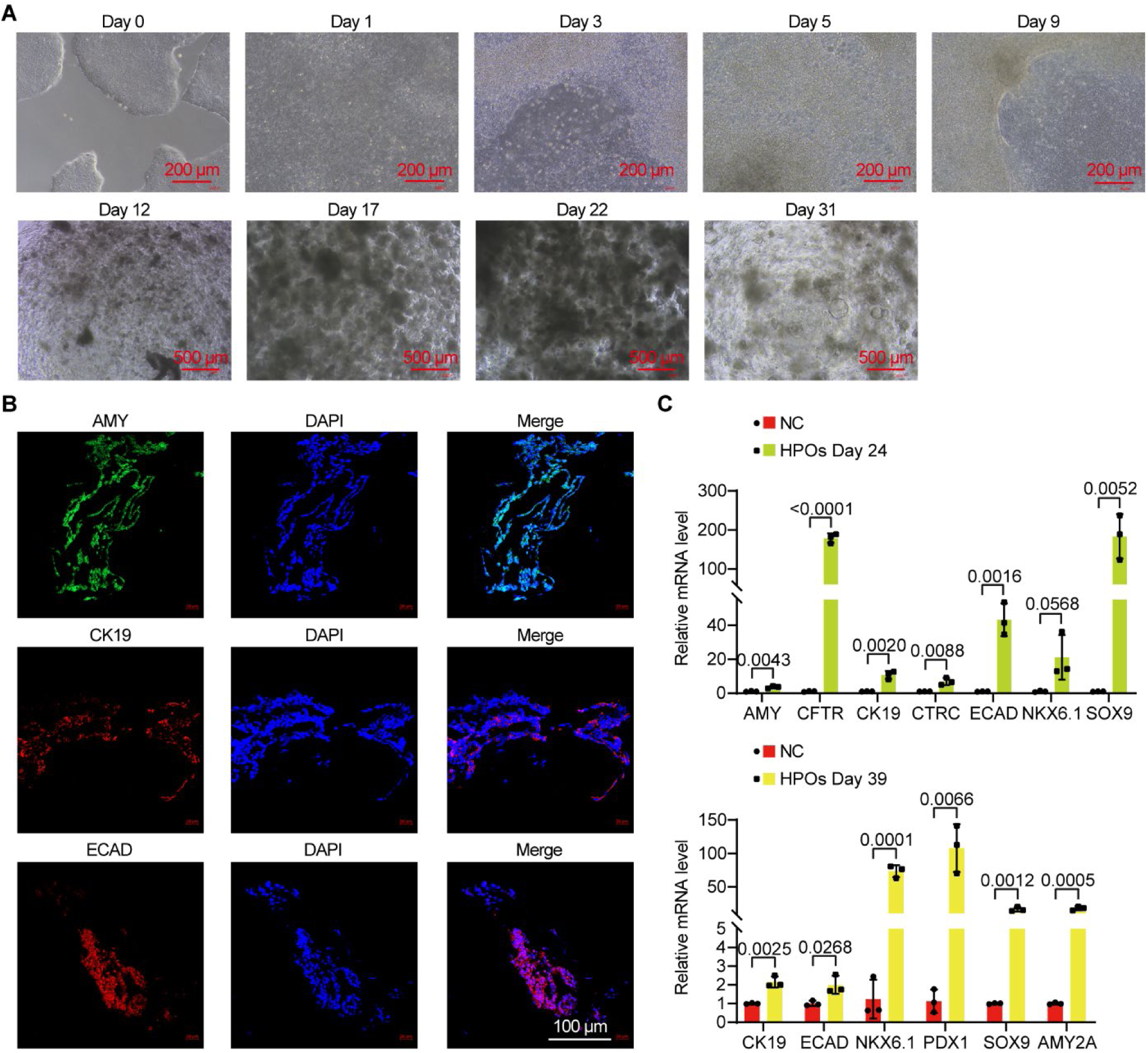
Characterization of HPOs. (A)Microscope images showing the morphological changes from hESCs to HPOs during differentiation. (B) IF data indicating the expression of AMY, CK19, and ECAD in differentiating HPOs. (C) qPCR results depicting mRNA levels of AMY, CFTR, CK19, CTRC, ECAD, NKX6.1, SOX9, PDX1, and AMY2A in differentiating HPOs at the indicated time points compared to undifferentiated hESCs.

## Appendix tables

**Table 1.** Antibodies used in Western blot analysis.

| **Antibody name** | **Catalog number** | **Manufacturers** | **Dilution** |
| --- | --- | --- | --- |
| ASC | 10500-1-AP | Proteintech, China | 1:500 |
| Pro-Caspase 1 | ab179515 | Abcam, UK | 1:500 |
| Cleaved-Caspase 1 | D57A2 | Cell Signaling<br>Technology | 1:500 |
| GSDMD | AF4012 | AFFINITY, China | 1:500 |
| Cleaved-GSDMD | ab215230 | Abcam | 1:500 |
| NLRP3 | 19771-1-AP | Proteintech | 1:500 |
| β-actin | 20536-1-AP | Proteintech | 1:1000 |
| CD9 | YT0782 | Immunoway, USA | 1:3000 |
| CD81 | YT5394 | Immunoway | 1:1000 |
| CD63 | YT0772 | Immunoway | 1:1000 |
| HRP-anti-Rabbit IgG | SA00001-2 | Proteintech | 1:10000 |

**Table 2.**
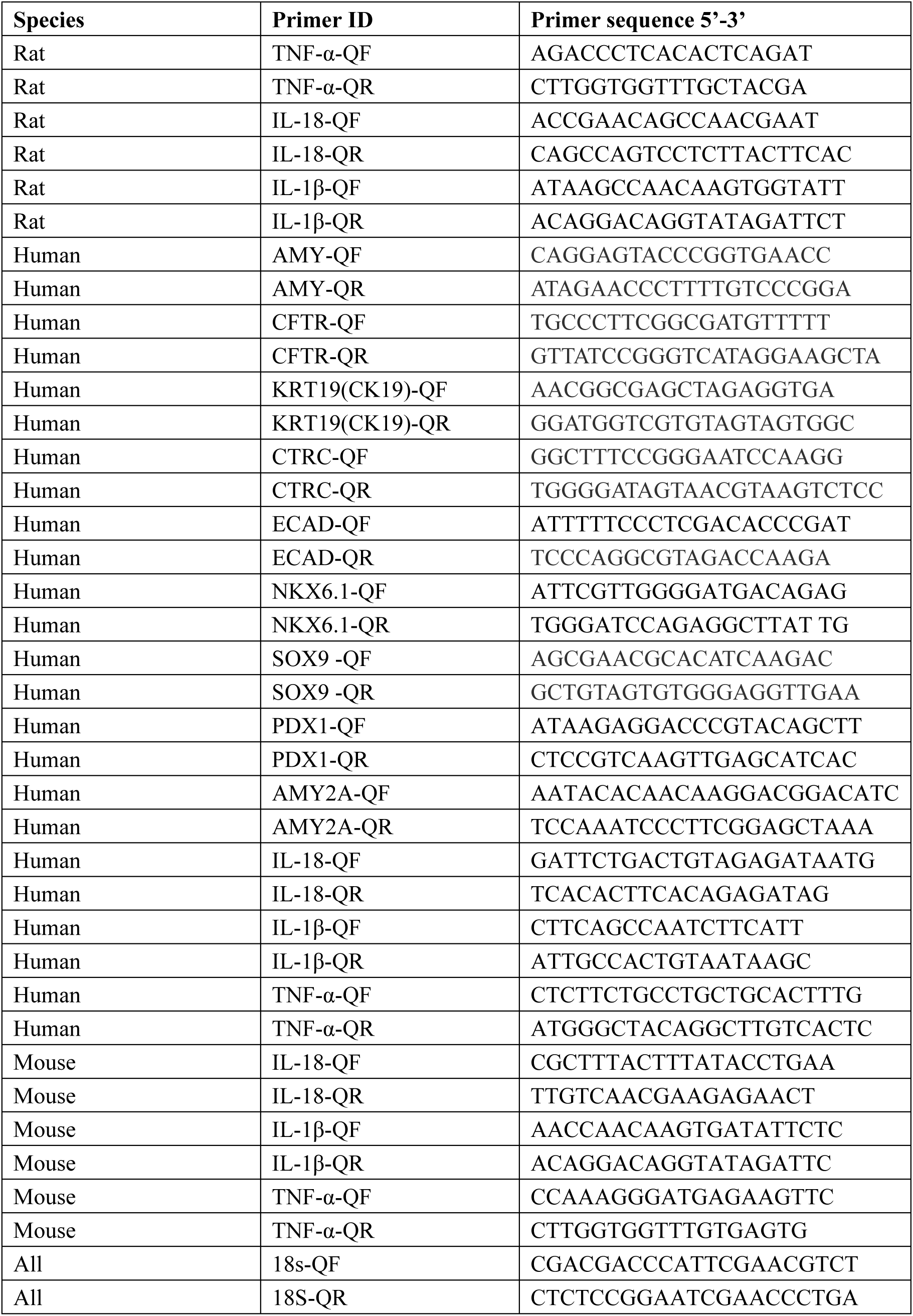
qPCR primers.

**Table 3.**
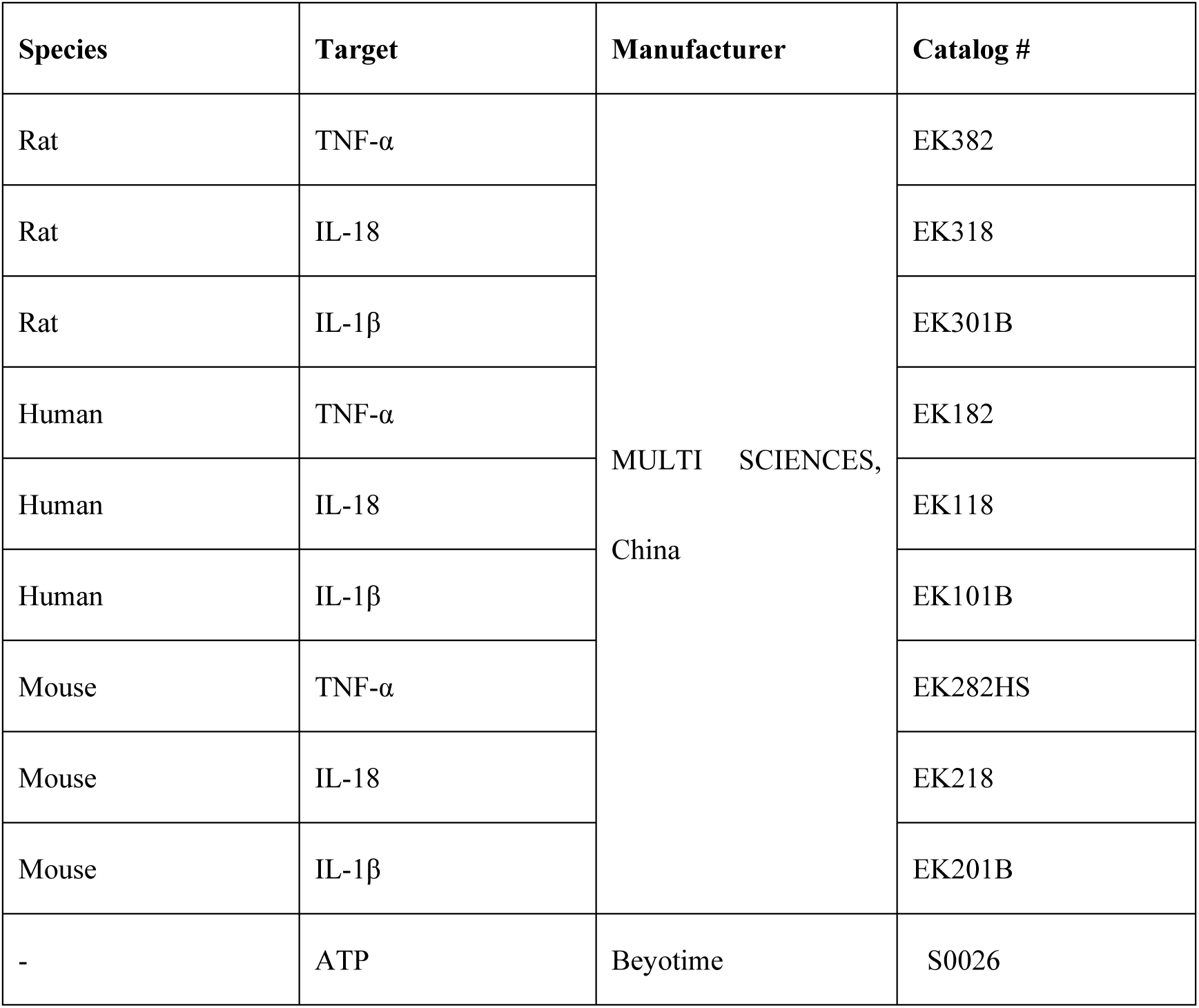
Kits for ELISA and ATP assays.

