## Supplemental Data 1 for "Emodin-Enhanced hUC-MSC Extracellular Vesicles Alleviate Acute Pancreatitis by Targeting Inflammation and Pyroptosis"

### Supplemental Figures

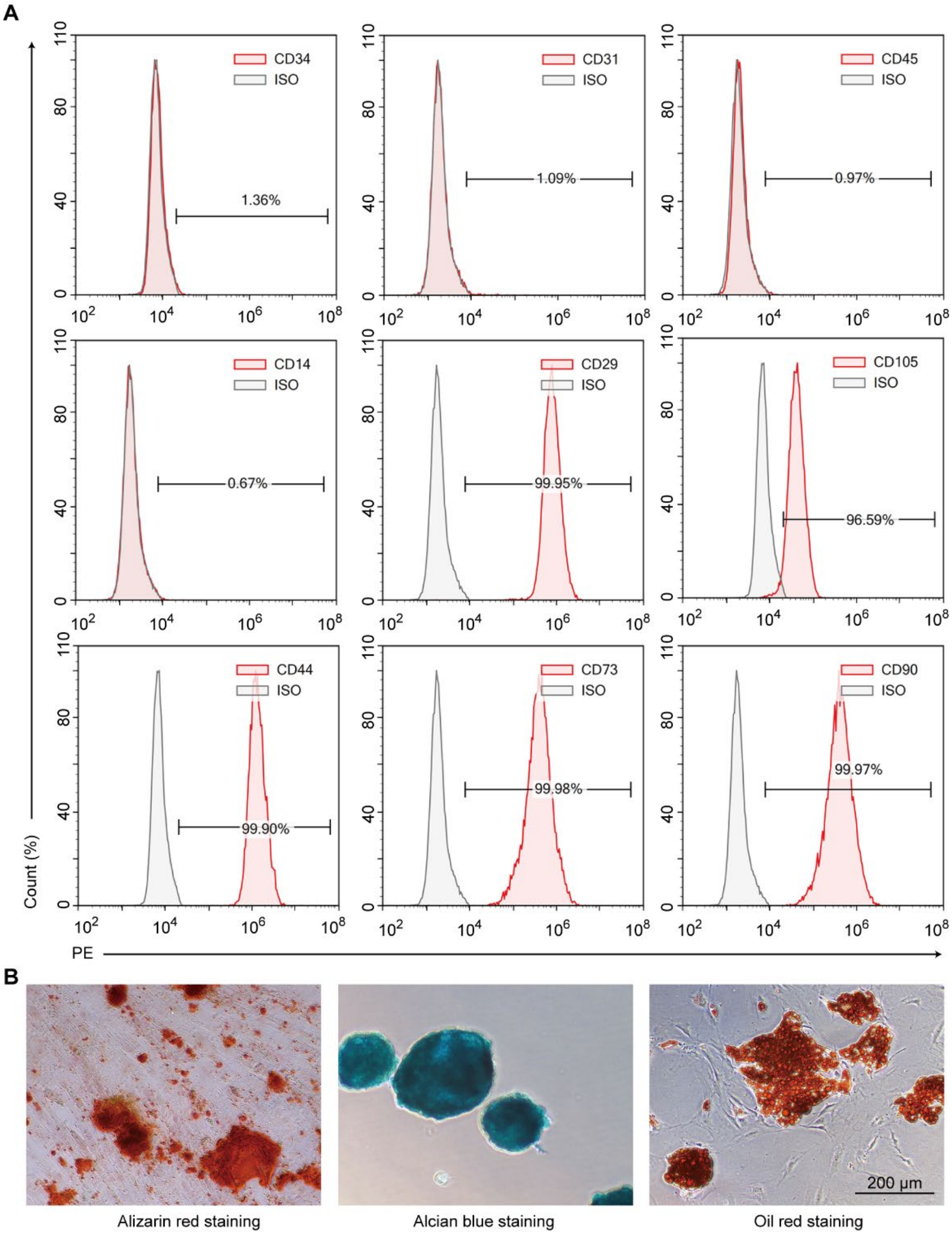


**Figure S1.** Characterization of hUC-MSCs. (A) Flow cytometry data showing negative expression of hematopoietic cell surface markers CD34, CD31, CD45, and CD14, and positive expression of MSC surface markers CD29, CD105, CD44, CD73, and CD90 on hUC-MSCs. (B) The effective differentiation of hUC-MSCs into osteocytes, chondrocytes, and adipocytes is confirmed by the positive staining results of Alizarin red, Alcian blue, and Oil red, respectively.

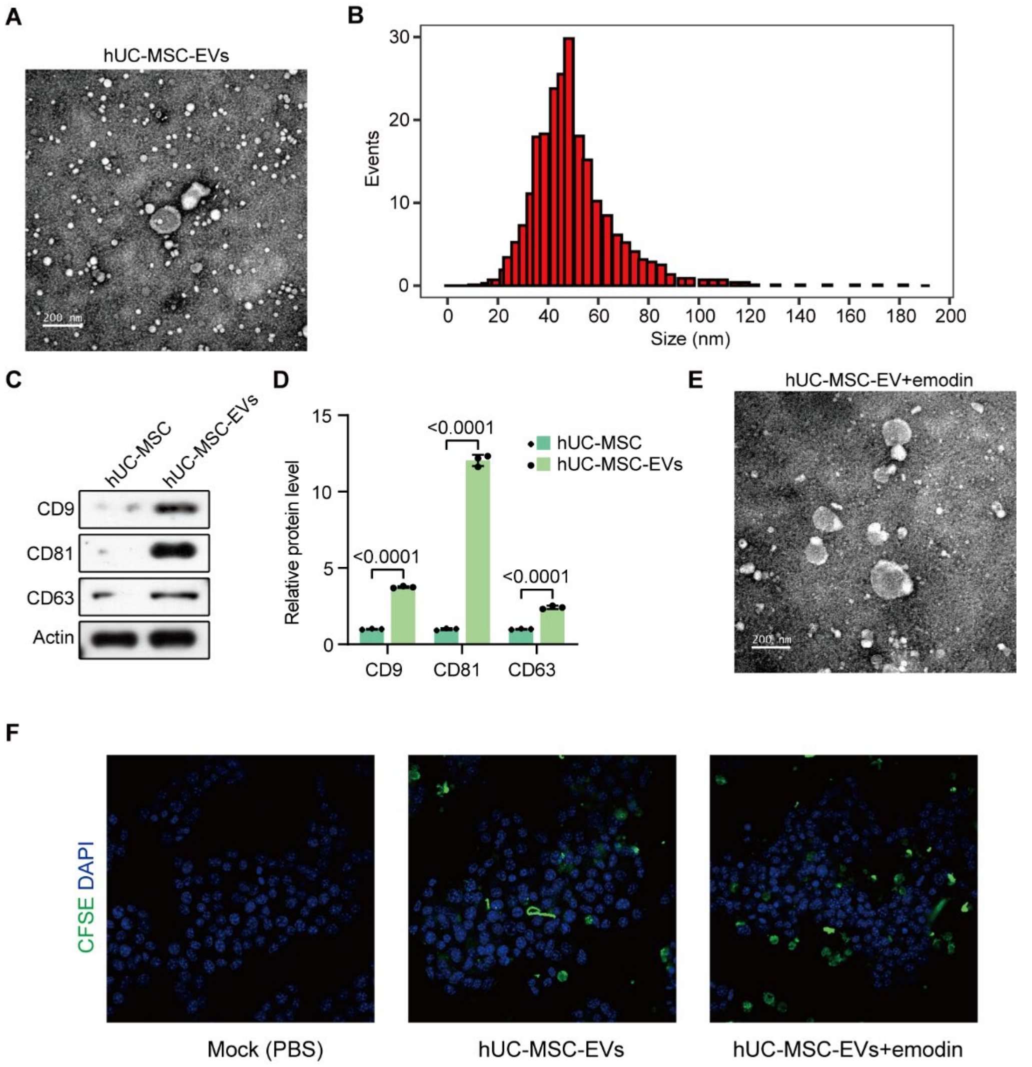


**Figure S2.** Identification of purified hUC-MSC-EVs. (A) TEM image showing the morphology of purified hUC-MSC-EVs. (B) NTA results depicting the size distribution of purified hUC-MSC-EVs. (C) Western blot data illustrating enrichment of CD9, CD81, and CD63 in purified hUC-MSC-EVs compared to hUC-MSCs. (D) Results of Western blot data analysis. (E) TEM image showing the morphology of emodin-loaded hUC-MSC-EVs. (F) Confocal images show the absorption of CFSE-labeled hUC-MSC-EVs and emodin-loaded hUC-MSC-EVs by AR42J cells.

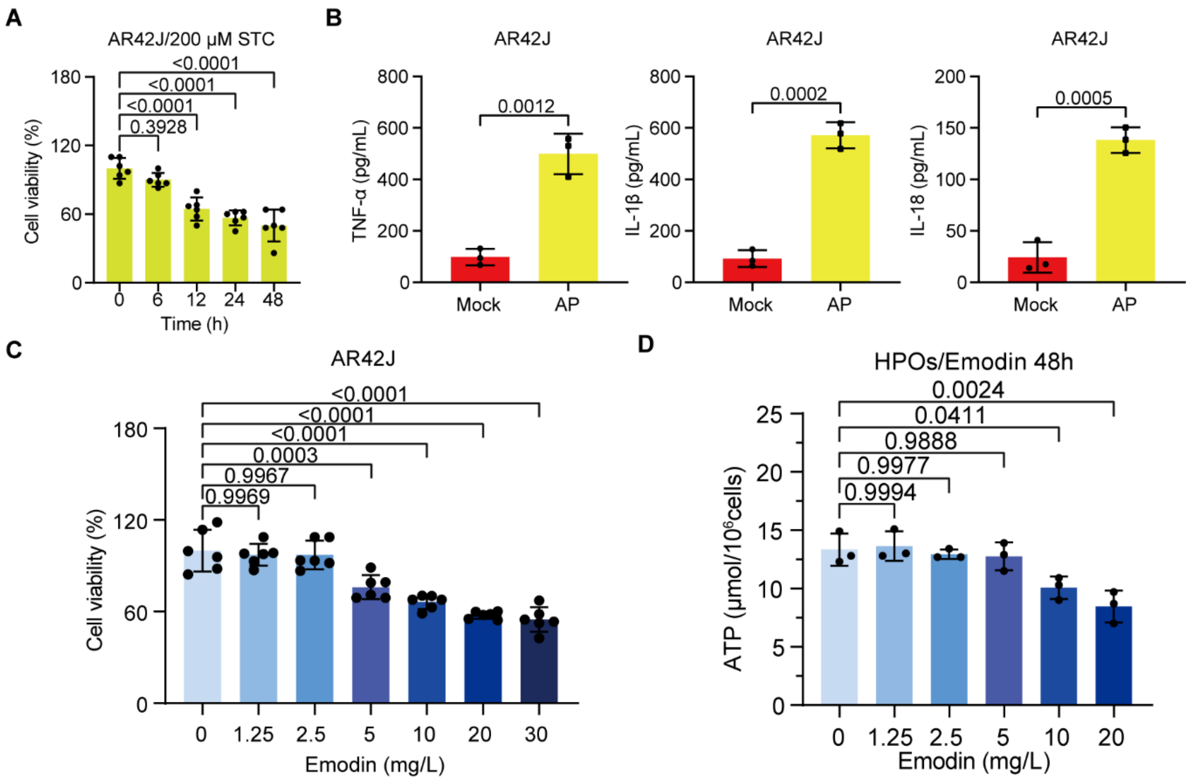


**Figure S3.** Optimization of STZ and emodin treatments. (A) MTT assay data showing the viability of cells exposed to 200 μM STZ for the indicated durations. (B) ELISA data indicating TNF-α, IL-1β, and IL-18 levels in the conditioned media of AR42J cells exposed to 200 μM STZ for 12 hours. (C) MTT assay results showing the viability of AR42J cells after treatment with specified concentrations of emodin. (D) ATP levels in HPOs following 48 hours of treatment with specified doses of emodin.

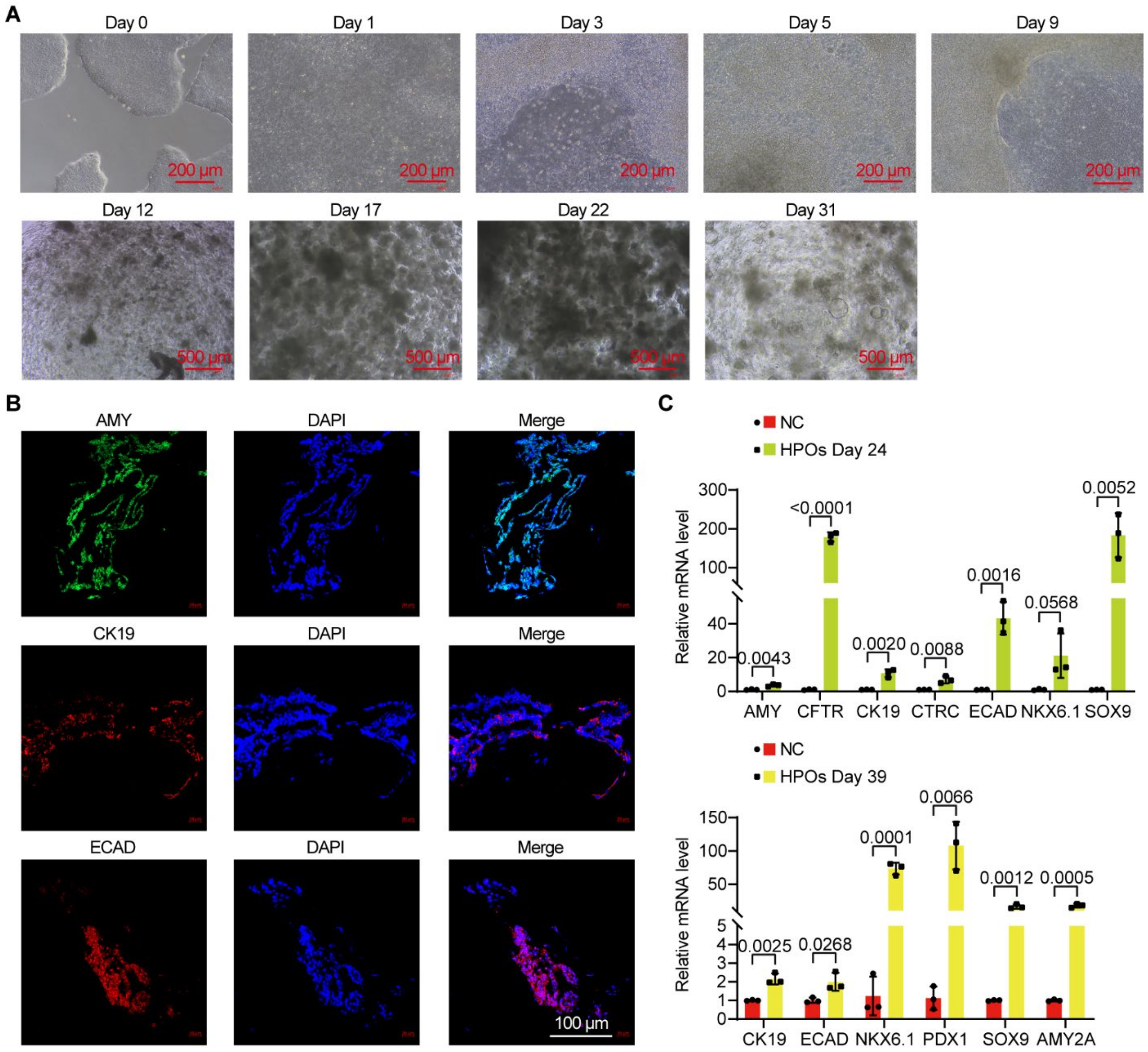


**Figure S4.** Characterization of HPOs. (A)Microscope images showing the morphological changes from hESCs to HPOs during differentiation. (B) IF data indicating the expression of AMY, CK19, and ECAD in differentiating HPOs. (C) qPCR results depicting mRNA levels of AMY, CFTR, CK19, CTRC, ECAD, NKX6.1, SOX9, PDX1, and AMY2A in differentiating HPOs at the indicated time points compared to undifferentiated hESCs.
